# The Role of Labile Iron on Brain Proteostasis; Could it be an Early Event of Neurodegenerative Disease?

**DOI:** 10.1101/2023.11.20.567981

**Authors:** Aiyarin Kittilukkana, Jannarong Intakhat, Chalermchai Pilapong

## Abstract

Iron deposits in the brain are a natural consequence of aging. Iron accumulation, especially in the form of labile iron, can trigger a cascade of adverse effects, eventually leading to neurodegeneration and cognitive decline. Aging also increases the dysfunction of cellular proteostasis. The question of whether iron alters proteostasis is now being pondered. Herein, we investigated the effect of ferric citrate, considered as labile iron, on various aspects of proteostasis of neuronal cell lines, and also established an animal model having a labile iron diet in order to evaluate proteostasis alteration in the brain along with behavioral effects. According to an *in vitro* study, labile iron was found to activate lysosome formation but inhibits lysosomal clearance function. Furthermore, the presence of labile iron can alter autophagic flux and can also induce the accumulation of protein aggregates. RNA-sequencing analysis further reveals the upregulation of various terms related to proteostasis along with neurodegenerative disease-related terms. According to an in vivo study, a labile iron-rich diet does not induce iron overload conditions and was not detrimental to the behavior of male Wistar rats. However, an iron-rich diet can promote iron accumulation in a region-dependent manner, particularly in the cortex. By staining for autophagic markers and misfolding proteins in the cerebral cortex, the iron-rich diet was actually found to alter autophagy and induce an accumulation of misfolding proteins. These findings emphasize the importance of labile iron on brain cell proteostasis, which could be implicated in developing of neurological diseases.

**Graphical abstract:** 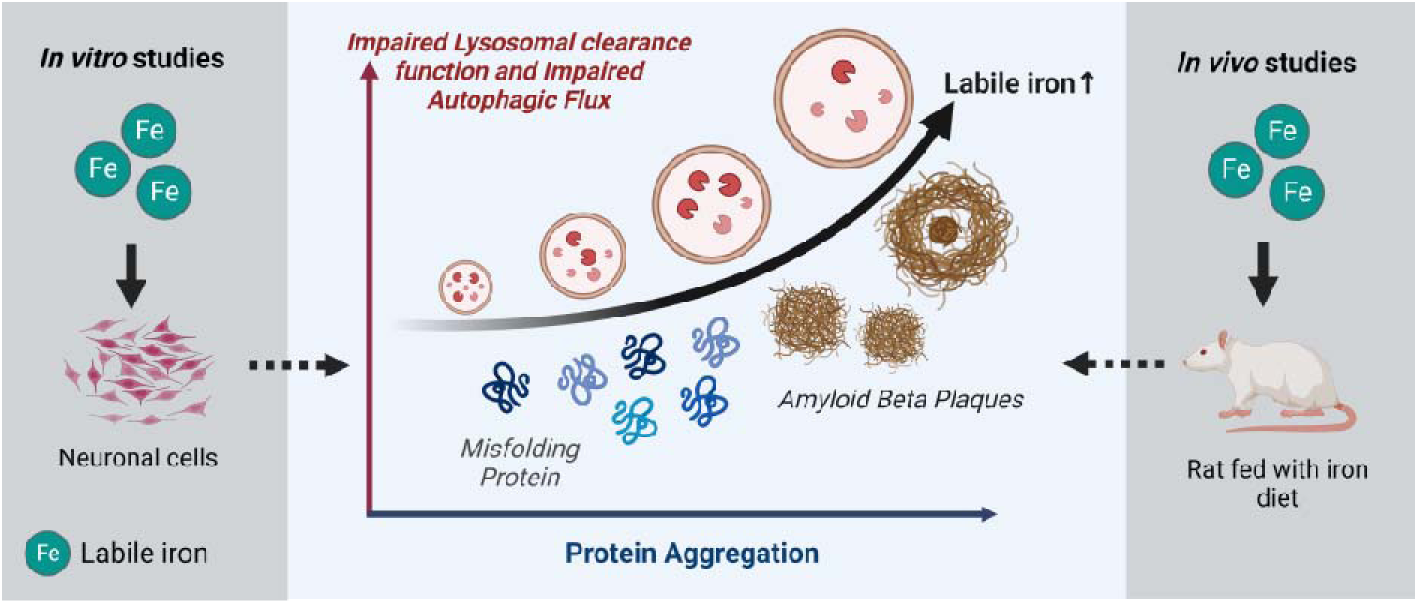

## Introductions

Iron has been known for decades as a vital element in the human body, helping to facilitate the creation of hemoglobin for oxygen transport, DNA synthesis, and ATP generation, as well as myelination, neurotransmitter formation, and synapse function in the brain (1, 2). In addition to ferritin and transferrin (Tf)-bound iron, the brain generally contains labile iron, an iron that binds to small ligands such as citrate and are highly active (3). The research has demonstrated that labile iron in the brain contributes to neuronal functioning (3, 4). The labile iron acts as both a static and dynamic biological function (5). In addition, it is evidenced that high levels of labile iron accelerates brain aging and cognitive decline (6). Generally, iron must be strictly controlled inside the brain; otherwise, it can produce oxidative stress via the Fenton reaction and cell death through ferroptosis can occur (7). There is accumulating evidence that iron could be one of the risk factors for neurodegenerative diseases (4, 8–10).

Proteostasis is a crucial process that regulates all aspects of protein within the cells. This process involves interconnection networks that influence synthesis to the degradation of protein (11). In the brain, sustaining this process is required for driving brain function properly. It is widely observed that the dysfunction of brain proteostasis is considered as a hallmark of aging, and multiple neurodegenerative diseases are characterized by dysfunctional proteostasis including lysosomal dysfunction and impaired cellular waste disposal system in the lysosomes and the ubiquitin-proteasome system (UPS) (11). Along with dysregulated proteostasis, as mentioned above, accumulation of iron is also one of the brains aging hallmarks. Whether iron can alter proteostasis is now being determined. Iron has been found to influence lysosome biology and function. Recently, a study by Xiao et al. revealed that iron inhibited autophagosome-lysosome fusion, hence it enhances the accumulation and transmission of alpha-synuclein. Moreover, iron-induced AKT/mTORC1 signaling inhibits the synthesis and nuclear translocation of nuclear transcription factor EB (TFEB), the main transcriptional regulator of lysosome biosynthesis and autophagosome-lysosome fusion (12). Furthermore, iron-rich lysosomes can promote lysosomal fragility and impairs lysosomal function through ROS-induced lysosomal membrane permeabilization. Lysosomal membrane rupture results in the release of cathepsins and other hydrolases from the lysosomal lumen into the cytoplasm (13). In addition, iron also can bind with amyloid-beta and tau protein inducing neurotoxicity (14, 15). Several articles have reported that amyloid beta has the ability to attach to iron and can convert iron (III) to iron (II) *in vitro* (16–18). The studies of Everett et al. suggested that amyloid beta can accumulate iron (III) within amyloid aggregates, resulting in the reduction of iron (III) to a redox-active iron (II) (19).

The aforementioned evidence indicates that labile iron could affect proteostasis, which could be crucial factor facilitating brain aging and could be implicated in developing neurological disorders later on. Currently, the connection between labile iron and different aspects of biology and function of proteostasis is still unclear. Herein, we aimed to investigate the role of labile iron in lysosomal biology and functions, autophagy flux, and the induction of misfolding protein accumulation. In this study, two models were utilized: cell culture and animal experimentation. In *in vitro* experiments, neuronal cell lines (SH-SY5Y) were used to examine the biological effects of labile iron. The *in vivo* experiments conducted on animals were designed to correlate with the *in vitro* study. The behavior test and tissue analysis of the proteostasis markers were conducted.

## Materials and Methods

### 1. *In vitro* experiments

#### Preparation of solutions of labile iron

Ferric citrate (LOBA CHEMI) was chosen as the labile iron representative for this study. The solution of ferric citrate (hereafter referred to as labile iron) was prepared by dissolving in DI water and then sonicated at room temperature until thoroughly dissolved. Next, the obtained labile iron solution was subjected to sterilization by filtering through a 0.22 μm cellulose acetate syringe filter membrane (Sartorius).

#### Cell Culture

Neuroblastoma Cell Line SH-SY5Y (Elabscience) (hereafter referred to as neuronal cells) was used in all *in vitro* models. The neuronal cells were cultured in DMEM HAM F-12 (Caisson Laboratories) supplemented with 15% Fetal Bovine Serum (FBS) (Carpricorn) and 1% penicillin–streptomycin solution (Caisson Laboratories) and maintained at 37 °C in a 5% CO_2_ atmosphere.

#### Cell Counting Assay

The cells were seeded in a 6-well plate (3 × 10^5^ cells per well) at 37 °C under a 5% CO_2_ atmosphere overnight. Next, the cells were incubated with 0, 10, 100, and 500 µM of labile iron for 24 and 48 hours. After that, the culture medium was discarded, and the cells were washed twice with the PBS buffer. The cells were detached by trypsinization, and the number of viable cells was counted using a flow cytometer (CytoFLEX).

#### Cell Death Assay

The cells were seeded in a 6-well plate (3 × 10^5^ cells per well) at 37 °C in a 5% CO_2_ atmosphere overnight. Next, the cells were incubated with 0, 10, 100, and 500 µM of the labile iron for 24 and 48 hours. After that, the culture medium was discarded, and the cells were washed twice with a Phosphate Buffer Saline (PBS) buffer. The cells were stained with Hoechst 33342 (APEXBIO) and Propidium iodide (PI) (United States Biological) at 37°C for different lengths of time. Afterward, the cells were observed under a fluorescent microscope (Nikon, Eclipse Ts2) with a blue channel for Hoechst and a red channel for PI. The cell death was assessed by comparing blue to red fluorescent signals in each sample.

#### Determination of intracellular iron levels

For quantitative analysis, the cells were seeded in a 6-well plate (3 × 10^5^ cells per well) at 37 °C in a 5% CO_2_ atmosphere overnight. Next, the cells were incubated with 0, 10, 100, and 500 µM of labile iron for 24 and 48 hours. After that, the culture medium was discarded, and the cells were washed twice with a PBS buffer. The attached cells were detached by trypsinization, and the cells were lysed with 6.5% Nitric acid (HNO_3_) and boiled for 10 minutes at 95°C. The lysed solutions were added with Potassium thiocyanate (KSCN) (Loba Chemie). The UV-VIS spectrophotometer (Perkin Elmer) was employed to measure absorbance spectra. The wavelengths were adjusted between 480 nm and 800 nm.

For qualitative analysis, the cells were seeded in a 6-well plate (3 × 10^5^ cells per well) at 37 °C in 5% CO2 atmosphere overnight. Then, the cells were incubated with 0, 10, 100, and 500 µM of labile iron for 24 hours. After that, the culture medium was discarded, and the cells were washed twice with the PBS buffer. Next, the samples were incubated with Prussian’s blue solution containing ferrocyanide acid solutions (5% Hydrochloric acid (HCl) and 5% potassium ferrocyanide (Merck)) at a 1:1 ratio for 30 minutes. After washing, the stained cells were observed under an inverted microscope (Nikon, Eclipse Ts2).

#### Cell Cycle Analysis

The cells were seeded in a 6-well plate (3 × 10^5^ cells per well) at 37 °C in a 5% CO_2_ atmosphere overnight. Next, the cells were incubated with 0, 10, 100, and 500 µM of labile iron for 24 and 48 hours. After washing, the cells were detached by trypsinization, and washed twice with PBS. The cells were then fixed with 70% cold ethanol at 4°C for 30 minutes. The fixed cells were washed twice with PBS, and further incubated with 5 μL of TritonX-100 (BioBasic) in PBS, 50 μL of 2 mg/mL RNase A (United States Biological), and 5 μL of 1 mg/mL PI at 37 °C for 30 minutes. The fluorescent intensity of the stained cells was measured with a flow cytometer. The analysis of the cell cycle was performed using the FlowJo v10.8.1

#### DNA damage Assay

Protein histone H2A.X and phosphorylated H2A.X (γH2AX) were performed according to the manufacturer’s instructions (Luminex). Briefly, the cells were seeded in a 6-well plate (3 × 10^5^ cells per well) at 37 °C in a 5% CO_2_ atmosphere overnight. Then, the cells were incubated with 0, 10, 100, and 500 µM of labile iron for 24 and 48 hours. The cells were collected and fixed with a fixation buffer. After that, the antibody cocktails were added to each sample and were incubated for 30 minutes at room temperature. The cells were then subjected to measure by flow cytometer. The analysis of DNA damage was performed using FlowJo v10.8.1.

#### Live Cell Imaging of lysosome

The cells were seeded in a 6-well plate (3 × 10^5^ cells per well) at 37°C in a 5% CO_2_ atmosphere overnight. Next, each sample was incubated with 0, 10, and 100 µM of labile iron for 24 and 48 hours. After washing, each sample was incubated with LysoTracker Red DND99 (Thermo Fisher Scientific) at 37°C for 30 minutes and were observed under a fluorescent microscope. For quantification assay, the cells were detached by trypsinization and incubated with the same probes. These cells were then measured by flow cytometer.

#### Neutral Red Retention Assay

To observe the lysosome by neutral red staining, the cells were seeded in a 6- well plate (3 × 10^5^ cells per well) at 37°C in a 5% CO_2_ atmosphere overnight. Next, the cells were incubated with 0, 10, and 100 µM of labile iron for 24 and 48 hours. After washing, the cells were stained with neutral red (May & Baker) (0.8 mM in final concentration) at 37°C for 30 minutes. Subsequently, the cells were washed twice with PBS and were supplemented with Hank’s Balanced Salt Solution (HBSS). Next, the stained cells were observed under a light microscope in a time-dependent manner.

To study the clearance function of the lysosome, the cells were seeded in a 24- well plate (5 × 10^4^ cells per well) at 37°C in a 5% CO_2_ atmosphere overnight. The cells were then incubated with 0 and 100 µM of labile for 24 and 48 hours. The cells were further incubated with a neutral red solution for 30 minutes in a fresh medium. Next, the cells were washed twice with PBS and were renewed with HBSS. Immediately, the stained cells were subjected to measurements of the absorbances of 455 nm and 525 nm at 37 °C using a plate reader equipped with kinetic mode (BioTek SynergyH4 Hybrid Reader).

#### Protein Aggregation Staining Assay

The cells were seeded in a 6-well plate (3 × 10^5^ cells per well) at 37°C in a 5% CO_2_ atmosphere overnight. Next, the cells were incubated with 0, 10, and 100 µM of labile iron with and without β-Amyloid Peptide (Aβ_1-42_) (rPeptide) for 24 and 48 hours. After washing, the cells were stained with Thioflavin T (Th T) (Abcam) (final concentration 20 µM) at 37°C for 1 hour. After that, the cells were washed and observed under a fluorescent microscope.

#### Immunofluorescent staining

The cells were seeded in a 6-well plate (3 × 10^5^ cells per well) at 37°C in a 5% CO_2_ atmosphere overnight. Later, the cells were incubated with 0, 10, and 100 µM of the labile iron for 24 and 48 hours. After washing, the cells were fixed in 4% paraformaldehyde (Sigma Aldrich) at 37°C for 5 minutes and were washed with PBS 3 times. Next, the cells were permeabilized in 0.1% TritonX 100 at room temperature for 15 minutes. After washing, the cells were incubated with anti-LC3B or anti-SQSTM1/P62 antibody (eLabscience) at 4°C overnight. Subsequently, the cells were further incubated with Dylight 488 conjugated anti-rabbit secondary antibody (Thermo Scientific) for 2 hours in the dark at room temperature and were washed with PBS 3 times. The cells were then observed under a fluorescent microscope.

#### RNA-Sequencing

The cells were seeded in a 6-well plate (3 × 10^5^ cells per well) at 37°C in a 5% CO_2_ overnight. Next, the cells were incubated with 0, 10, and 100 µM of labile iron for 24 hours. The detached cells were used for RNA isolation using NucleoSpin RNA Plus kit (MN) in accordance with the manufacturer’s instructions. The quality, integrity, and purity of total RNA were checked using an Agilent Bioanalyzer 2100 system, agarose gel electrophoresis, and a Nanodrop spectrophotometer. After a quality control (QC) procedure was performed, messenger RNA (mRNA) was purified using poly-T oligo-attached magnetic beads. Complementary DNA (cDNA) libraries were constructed, according to the manufacturer’s recommendations (Novogene Corporation, Beijing, China). All libraries were sequenced using an Illumina HiSeq PE150 platform. The library construction and sequencing were performed by the Novogene Corporation. After receiving the raw data, QC check was performed by FastQC and adapter sequences were removed by Cutadapt. The data was then aligned with HG38 Human genome by the STAR. Quantification of genes was performed by FeatureCounts. Transcriptome assembly and differential expression analysis was performed by Cufflinks. The gene enrichment tools from the Nucleic Acid SeQuence Analysis Resource (NASQAR) were utilized for functional enrichments such as Kyoto Encyclopedia of Genes and Genomes (KEGG pathway) and Gene ontology (GO).

### 2. *In vivo* experiments

#### Animal use protocol

The adult male 8-week-old Wistar rats were purchased from Nomura Siam International Co., Ltd. Bangkok, Thailand. All animals were maintained in a pathogen-free husbandry with controlled ventilation and temperature. Two rats were housed in each cage. The rats were raised in the Standard Laboratory Animal Center, Chiang Mai University conditions with a 12-hour light-dark cycle (lights on from 06:00 to 18:00) at 21°C with free access to food and water ad libitum. All procedures involving animal care were processed in accordance with the regulations of IACUC - American Association for Laboratory Animal Science and were approved by the Animal Care and Use Committee at the Laboratory Animal Center, Chiang Mai University (Protocol number: RT014-2562).

The experimental rats were divided into two groups, normal diet (ND, n = 6) and labile iron diet (ID, n = 6). The rats in ND group were given a commercial rodent diet but the rats in ID group were given a commercial rodent diet containing ferric citrate (0.2% iron) for 8 weeks (called the “induction phase”). At week 9, all rats were continuously housed without any treatment and were given a commercial rodent diet for another 8 weeks (called the “satellite phase”). At the end of the experiment, all rats were sacrificed, and brains were collected for further analysis.

#### Behavioral experiments

The Behavioral experiments were performed during the same circadian period (9.00 to 15.00) at the Laboratory Animal Center, Chiang Mai University. Behavior experiments were performed at the end of the induction phase and satellite phase.

The Morris water maze (MWM) test has been used to study spatial learning and memory, and the experiment was performed according to previous protocol (20). Briefly, the test consisted of a circular swimming pool filled with opaque water and a platform submerged in the water. In the training phase, the rats were allowed to freely swim until they found the submerged platform to escape from the water within 120 seconds. Each rat was trained four rounds per day. The time at which the rat reached the platform in each trial was called training latency time, which was considered a measure of the acquisition of spatial navigation abilities. In the testing phase (probe trial), the platform was removed from the pool, and the rats were allowed to swim again. Time at which the rat spent in a quadrant of the pool that previously had the platform was recorded and this time was used to assess spatial memory.

The Radial arm maze (RAM) test has been used to study spatial learning as well as working and reference memory, and the experiment was performed according to this protocol (21). Briefly, the eight-arm maze consisted of a central platform designed to assess the ability of animals to recall which arms provided rewards (food) and which did not. The spatial working memory test was conducted with four arms held open containing rewards and four arms that were closed. The rats should visit each arm only once. The rats that visited open arms more than once counted one score as a working memory error. The spatial reference memory test was conducted with all eight arms open, and the reward contained only four arms that were previously closed in the spatial working memory test. The rats that visited with no reward arm counts one score as a reference memory error.

The Novel object recognition (NOR) test has been used to determine the learning and memorization of familiarity with objects that have been seen or experienced at a previous time. The experiment was performed according to this protocol (22). Briefly, the test was divided into two parts. In the first part, two objects having the same shape, size, and color were placed in the designed box. In the second part, one of the previous objects and a novel (new) object were both placed in a designed box. The test assessment was done by recording the time that it took the rat to investigate each object. Afterward, the memory index (Recognition Index) was calculated with equation 1.

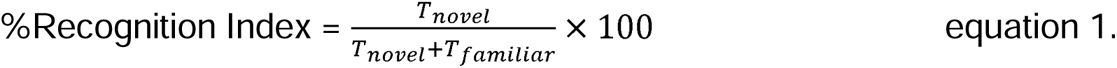

T_novel_ = time that the rat spent to observe new object

T_familiar_ = time that the rat spent to observe familiar object

#### The detection of labile iron in Magnetic Resonance Imaging (MRI) *In vivo* model

The MRI scans were performed at the end of the induction and satellite phases. All rats were anaesthetized using a 40 mg kg^-1^ body weight of thiopental via intraperitoneal injection and the brain was scanned by an MRI scanner (Philips Ingenia 1.5 T) using a T_2_*-weighted gradient echo pulse sequence technique. Echo time (TE) was set as 1-32 milliseconds, repetition time (TR) was set as 100 milliseconds using a head and neck coil for receiving the signal. After scanning, the images were selected to determine the region of interest (ROI), and signal intensity was measured using Philip DICOM viewer R3.0 software. The T_2_* values were determined by curve fitting with equation 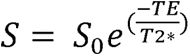 The transverse relaxation rate (*R*_2_*) was also determined with equation R2* = 1/T2*

#### Tissue iron determination

The quantification of tissue iron was performed according to a previous study (23). Briefly, the brain tissues were homogenized using a Dounce homogenizer. The homogenate tissues were then centrifuged at 15000 rpm at 4°C for 30 minutes. Next, the supernatants were collected and were mixed with Trichloroacetic acid (TCA) (Sigma) and HCl solution at a ratio of 1:1. The samples were boiled at 95°C for 1 hour. After that, the samples were centrifuged at 14000 rpm for 30 minutes. The supernatant was then collected and was well mixed with 69% nitric acid (HNO_3_) for 30 minutes. The KSCN was then added to the supernatant. The iron content was identified by a UV-VIS spectrophotometer.

For iron staining, the tissue slides were deparaffinized with two changes of xylene (RCI Labscan) for 5 minutes each. Liquid was removed and slides were subsequently rehydrated in two changes of absolute ethanol at 90%, and 80% ethanol for each. The slides were rinsed in tap water and were place in a PBS wash jar for further rehydration. The slides were incubated for 30 minutes in Prussian’s blue solution containing ferrocyanide acid solution (5% HCl and 5% potassium ferrocyanide at a ratio of 1:1). After that, they were immediately rinsed with both tap water and deionized water. Subsequently, the slides were incubated with 0.05% 3,3′-Diaminobenzidine (DAB) (Sigma) solution for a few minutes, followed by a rinsing in PBS. The slides were dehydrated with ethanol and immersed again in xylene. The slides are mounted with Permount (Fisher Chemical). The slides are then examined and observed using a fluorescent microscope.

#### Immunohistochemistry assay

An immunohistochemistry experiment was used for determining autophagy markers. After deparaffinization and rehydration of the brain slides was completed, antigen retrieval was performed using a 10 mM sodium citrate buffer at 65°C for 10 minutes. To inactivate endogenous peroxidase, the slides were blocked with 0.3% H_2_O_2_ (Merck) and then the slides were permeabilized and blocked using 0.4% TritonX-100 and 5% Bovine Serum Albumin (BSA) (Capricorn) in PBS at room temperature. After that, the slides were incubated with an anti-LC3B antibody at 4 °C overnight. Afterward, the slides were further incubated with anti-rabbit secondary antibody (Jackson ImmunoResearch Laboratories, Inc) for 2 hours at room temperature. The slides were then washed and incubated with 0.05% DAB in 0.015% H_2_O_2_ in PBS for a few minutes. Finally, the slides were dehydrated and mounted with Permount. Analysis and observation of the slides was then done under a microscope.

#### Protein Aggregation staining

To evaluate the protein aggregation, Congo red (Biobasic) staining (24, 25) and Thioflavin T (ThT) staining (26) were performed. Briefly, after deparaffinization and rehydration of the brain slides, the slides were stained with 1% Congo red solution prepared in DI water for 15 minutes. After washing with tap water, the slides were differentiated quickly in an alkaline alcohol solution (1 ml of 1% NaOH and 100 ml of 50% ethanol) and were rinsed in tap water. After that, the slides were then dehydrated and mounted with Permount. Analysis and observation of the slides were done under a fluorescent microscope. The excitation of Congo Red is between 490 nm to 512 nm (27) and the emission of Congo Red is between 525 to 625_nm (28).

For ThT staining, the slides were stained with 1% ThT in DI water according to the same procedure as the Congo Red staining. There was no need to differentiate in alkaline alcohol solution. The excitation of ThT is from 385 nm to 450 nm and the emission is from 445 nm to 482 nm (29).

### 3. Statistical Analysis

Statistical analysis was performed using GraphPad Prism 8.0. All data are expressed as the mean ± standard deviation (SD) unless stated otherwise. Description of statistical analysis of different results is provided in the corresponding figure legends.

## Results and Discussions

In the present study, we selected ferric citrate for our investigation because it was considered to mimic the labile iron found in the brain (30, 31). We utilized neuronal cells (SH-SY5Y cell line) as a cellular model for *in vitro* investigation. It is well-established that excessive labile iron can be detrimental to brain cells. Therefore, biological evaluations associated with cytotoxic effects of labile iron were carried out. Firstly, the cell death analysis by Hoechst/PI assay was performed. Labile iron was found to induce cell death in a concentration and time-dependent manner (Figure 1A). The cells treated with 100 and 500 µM of labile iron showed a significantly higher percentage of cell death at both 24 and 48 hours. However, those of 10 µM treated had significant results only at 48 hours. of incubation. By assessing cell viability using a cell counting assay (Figure 1B), it was found that the labile iron could reduce the number of viable cells after 24 hours of incubation regardless of the dose of treatment. However, there was a significant change in percent cell count in 100 and 500 µM treated cells at 48 hours of incubation. There is no doubt that iron plays a role in cell cycle progression and can be harmful to the DNA via initiation of DNA damage. Thus, we checked whether labile iron is capable of disrupting cell cycle progression and initiating DNA damage or not. According to cell cycle analysis, it was found that the labile iron did not dramatically affect cell cycle progression (Figure 1C). As for determining any DNA damage, the results showed that only 500 µM of labile iron was found to increase total protein histone (H2AX), as compared to the control (Figure 1D). This is in good accordance with cytotoxicity tests where this dose exhibited a high toxicity against the cells. To determine the iron content in the cells, we performed quantitative analysis of intracellular iron content. As expected, iron was found to accumulate within the cells in a concentration-dependent manner (Figure 1E). Likewise, iron staining by Prussian’s blue assay also confirmed the existence of iron within the cells (Figure 1F). RNA sequence analysis suggested that the labile iron could activate cellular iron homeostasis (Figure 1G). From the above results, it was found that increased labile iron concentrations show a greater susceptibility to increases in the percentage of cell death, and H2AX. Nevertheless, the labile iron concentrations of 10 and 100 µM exhibit lesser toxicity to the cells. Consequently, we chose these dosages for subsequent investigation regarding proteostasis.

**Figure 1.**
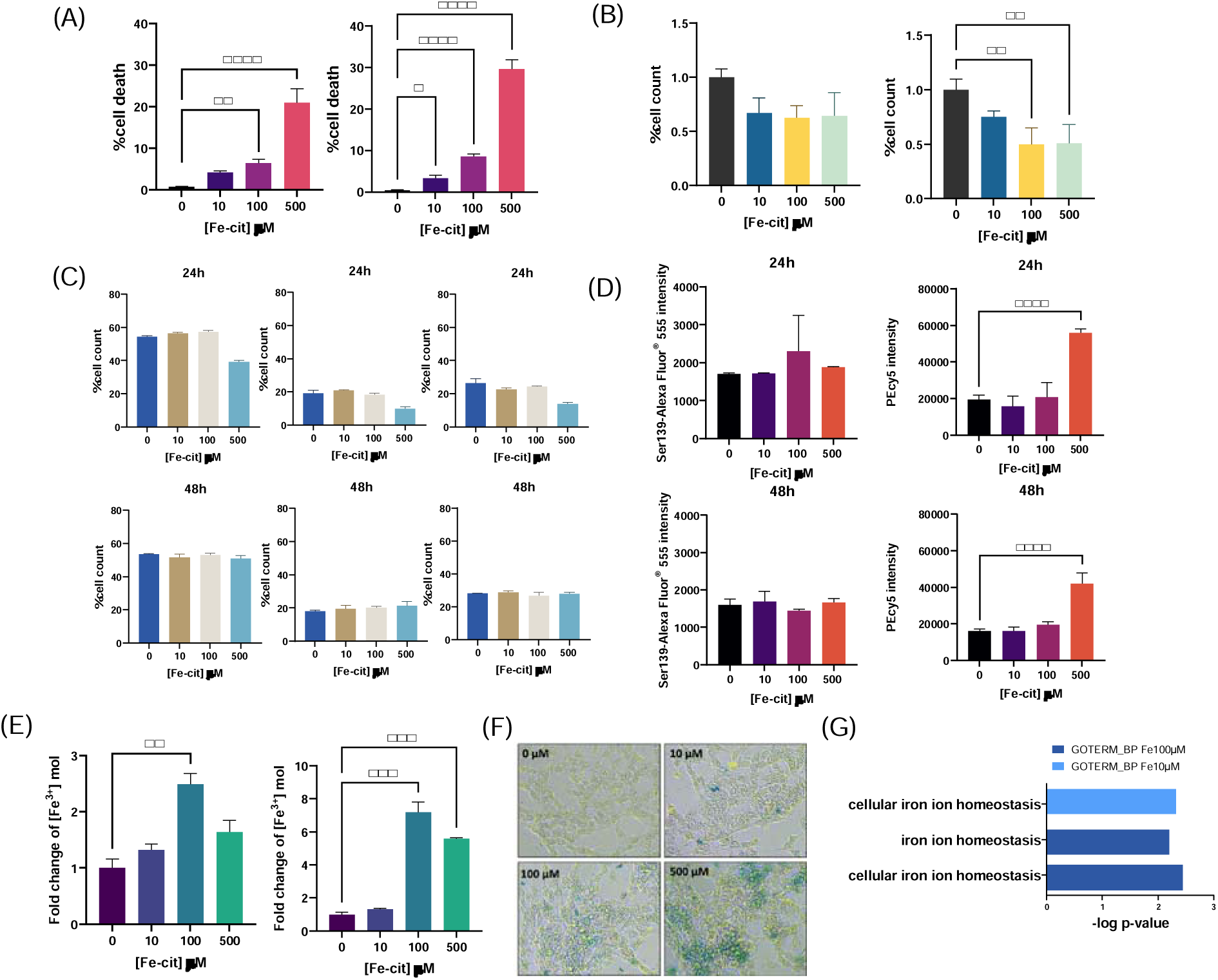
The effect of labile iron in SH-SY5Y cell lines. (A) Percentage of cell death was quantified by Hoechst/PI staining after incubating with labile iron for 24 hours (left) and 48 hours (right) (for 24 hours; 0, 10, 500 µM n=3 and 100 µM n=6) (for 48 hours.; 0, 500 µM n=3 and 10, 100 µM n=4). (B) Quantification of cell count percentage was assessed by using flow cytometry analysis after incubating with labile iron for 24 hours (left) and 48 hours (right) (n=3). (C) Cell cycle analysis was conducted to assess the distribution of cells across various phases; G1-phase (left), S-phase (middle), and G2/M-phase (right) (n=3). (D) Evaluation of DNA damage is conducted by quantifying the levels of total protein histone (H2AX) (PECy5) and histone H2AX phosphorylation (γH2AX) (Ser139-Alexa Fluor^®^ 555) (n=3). (E) Measurement of intracellular iron concentrations following 24 hours (left) and 48 hours (right) exposure to labile iron (n=2). (F) Iron detection was performed by Prussian’s blue staining method following 24 hours of exposure to labile iron. (G) An RNA-sequencing analysis was conducted to identify gene ontology (GO) terms that were differentially expressed and associated with biological processes in the cells treated with labile iron concentrations of 10 µM and 100 µM, with respect to untreated cells. * *p*<0.05, ** *p*<0.01, *** *p*<0.001, **** *p*<0.0001, by one-way ANOVA with Tukey’s multiple comparison test.

According to a previous study, lysosome was considered as the master regulator of iron (32). Thus, we aimed to study the effect of labile iron on lysosomes in the neuronal cells. We first investigated the lysosome mass and signal after incubation with the labile iron for different lengths of time. As observed by live cell imaging of lysosome by LysoTracker staining (Figure 2A) and neutral red staining (Figure 2B), it was found that the lysosomal signal had increased in a concentration and time-dependent manner. The quantification of lysosomal signal by flow cytometry showed that 10 µM of labile iron can significantly increase lysosome content but its content was found to reduce as the concentrations of the labile iron increased in both 24 and 48 hours of incubation (Figure 2C). As far as assessment of lysosome function was concerned, we employed neutral red (NR) retention assay to determine clearance function of lysosome. This assay was based on colorimetric analysis of NR that had accumulated in lysosomes of the cells (33). At a desired time, the cells were subjected to observe the red puncta (NR in lysosome) under an inverted microscope. As a result, the red puncta are clearly observed (denoted as 0 min) immediately after adding HBSS. As time passed, the red signal decreased, implying that the NR had cleared from the lysosomes (Figure 2D and 2E). The clearance kinetics of NR in lysosomes of cells treated with labile iron were examined using a microplate reader. The span and rate constant of clearance (*k*) were measured (as explained in Figure S1). We took advantage of span and *k* as parameters for assessing lysosome clearance function because they represented the amount of NR accumulated in lysosome and clearance kinetic of NR, respectively. As the results show in Figure 2F, the span and *k* values were found to decrease when the cells were incubated with the labile iron in a concentration-dependent manner, indicating that the labile iron could alter NR retention in lysosomes by lowering clearance rates. This result suggests that the labile iron could interrupt the clearance function of lysosomes.

**Figure 2.**
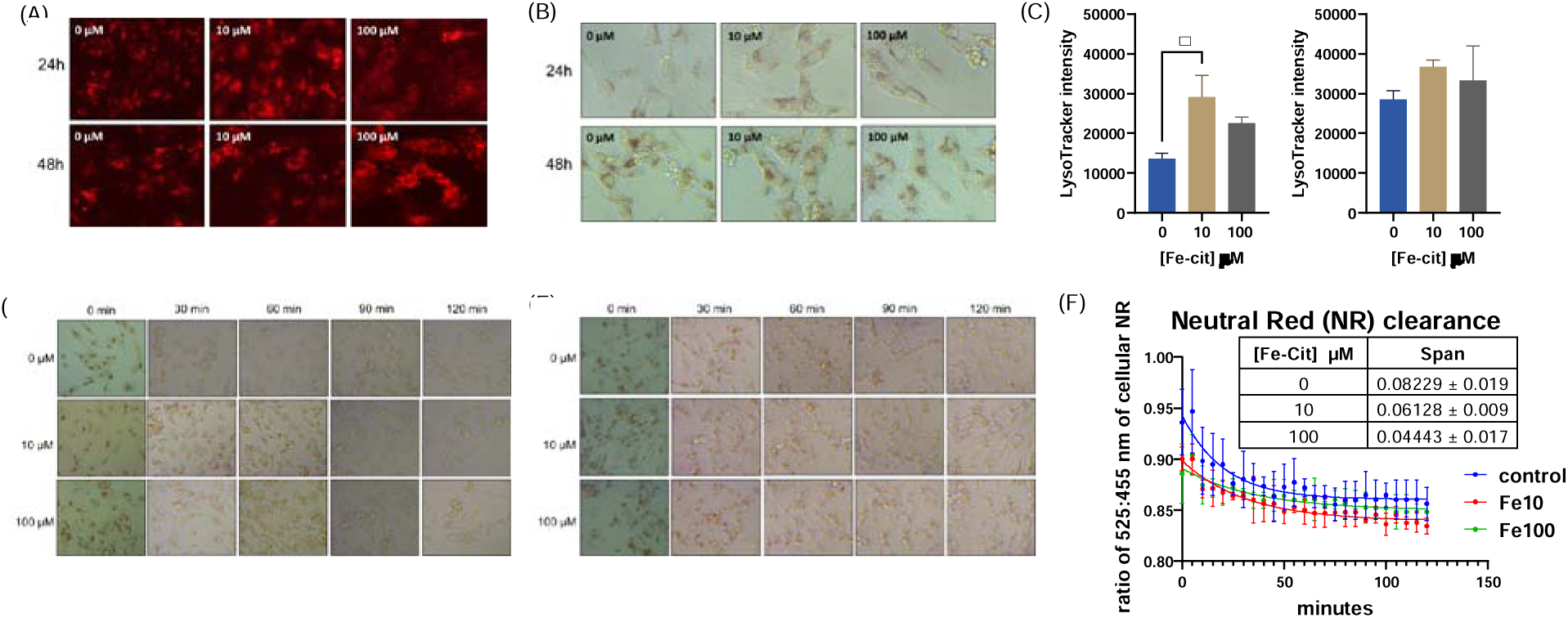
Labile iron impairs the mass and function of lysosomes in SH-SY5Y cell lines, hindering their ability for clearance. The visualization of lysosomes using LysoTracker staining (A) and neutral red staining (B). (C) Measurement of lysosomal mass by flow cytometry (n=3). Evaluation of the lysosomal clearance function by employing neutral red staining after subjecting to iron exposure for 24 hours (D) and 48 hours (E). (F) Exponential decay graph demonstrating the neutral red clearance rate and the span and rate at which neutral red is eliminated under different experimental conditions (n=3). Statistical analysis employed the one-way ANOVA. significance levels: * *p*<0.05 by one-way ANOVA with Tukey’s multiple comparison test.

Since labile iron exposure has been shown to alter lysosomal mass and signal, and lysosome function is closely related to autophagy process, we further investigated the expression of autophagy markers as a means of examining the cellular process of removing and recycling unnecessary or damaged components (34). In a typical manner, immunofluorescence assay of LC3B and p62 was carried out to assess autophagy marker and autophagic flux. According to LC3B and p62 staining (Figure 3A and 3B, respectively), labile iron was found to induce formation of LC3B and p62 puncta in both a concentration and dose-dependent manner. Normally, the presence of LC3B puncta on autophagosome membranes can be found during autophagy. However, p62, a protein that specifically targets ubiquitinated payloads for destruction by the autophagosome, has to be degraded (35). According to our results, LC3B was activated but p62 was not degraded, suggesting inhibition of autophagy flux mediated by labile iron. Based on the aberrant autophagic flux mediated by the labile iron, we determined whether labile iron could induce endogenous protein aggregation within the cells by using Thioflavin T (Th T) staining assay. The results showed that the protein aggregation was increased in both a concentration and time-dependent manners (Figure 3C). Similar results were observed when the cells were stimulated with misfolding peptides (Aβ_1-42_), labile iron was found to induce peptide aggregation (Figure 3D). It is known that many neurological disorders, including Alzheimer’s, Parkinson’s, and Huntington’s, are characterized by protein aggregation (36), thus induction of peptide and protein aggregation by labile iron could initiate early onset of pathological conditions related to such diseases, which can further disrupt cellular homeostasis and damage cells. It has been suggested that labile iron is one of the initiators of protein aggregation (14, 37) and the evidence presented herein promotes a similar manner of actions. Next, we checked transcriptomic profiles associated with proteostasis by GO and KEGG enrichments. Here we found that labile iron could be up-regulated various terms associated with ribosomes, translation processes, and ubiquitin activity in a dose-dependent manner (Figure 3E and 3F). Notably, there was a substantial increase in the enrichment score of lysosome-related and translation terms along with its negative regulation found in 100 µM treated cells suggesting that labile iron could somewhat alter the translation process. Such alteration has been shown to be associated with protein misfolding. Possibly, high expression of the translation mediated by labile iron could induce an over translation rate, resulting in less accurate translation which is prone to promote protein misfolding (38, 39). On the other hand, labile iron itself was capable of binding with amino acid in peptide chains, resulting in abnormal folding peptides, which could further aggregate to misfolding peptides or proteins (14, 15). As a consequence, a protein quality control system arrives to correct this abnormal peptide or protein misfolding, as observed in high expression of negative regulation of translation, especially, in 100 µM. To determine whether labile iron could induce an expression of gene associated to neurodegenerative diseases, RNA-seq analysis was conducted and various KEGG terms associated with neurodegenerative diseases were enriched (Figure S2). From the above observations, it can be inferred that proteostasis could be considered as a target for the labile iron, a high accumulation within the cells could induce proteostasis alteration, which is implicated in the development of neurological disorders.

**Figure 3.**
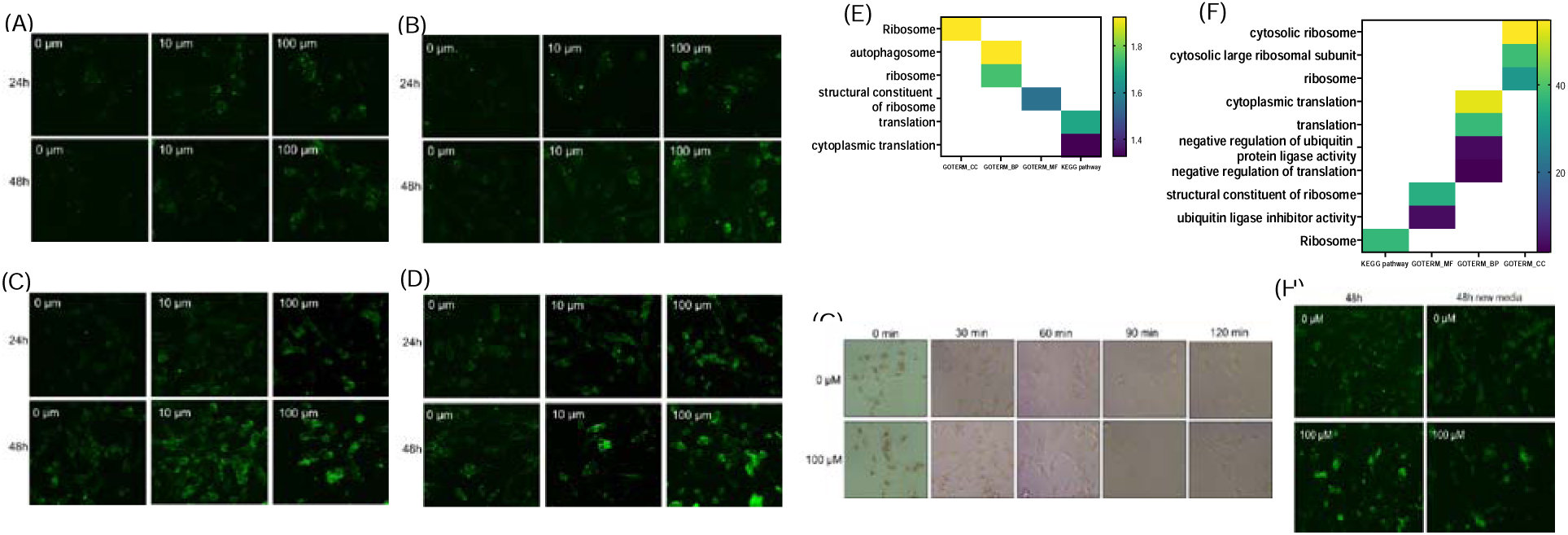
Labile iron has been found to affect proteostasis at both the cellular and transcriptomic levels in SH-SH5Y cell lines. The efficacy of media renewal in restoring lysosomal clearance function and accumulation of protein accumulation in SH-SY5Y cell lines. The autophagy marker LC3B (A) and p62 (B) were analyzed by immunofluorescent staining. The misfolding proteins were assessed by Th T staining under two conditions: without Aβ induction (C) and with Aβ induction (D), both under labile iron exposure conditions. RNA-sequencing analysis showed that the terms related to proteostasis were up-regulated after exposure with labile iron at concentrations of 10 µM (E) and 100 µM (F). (G) Lysosomal clearance function was assessed using a neutral red retention assay after 24 hours exposure to labile iron and the new cell culture medium without labile iron was then replaced before continuing the culture for an additional 24 hours (called “48 hours new media”). (H) The comparison of Th T staining between after prolonged exposure labile iron at 48 hours and after 48 hours after having changed the new media.

The above results indicate that labile iron has an effect on various aspects of proteostasis in neuronal cells including lysosome function, protein translation, and protein folding. Nevertheless, our quality-control system of cells enables the prevention of some biological processes linked to the onset of neurological disorders (40–42). Following this, we aimed to determine whether the disruption of proteostasis caused by labile iron can be done naturally or is easily alleviated. In a typical manner, the neuronal cells were treated with a culture medium containing 100 µM of labile iron for 24 hours, and then the fresh medium was renewed and was further incubated for another 24 hours. Finally, lysosome clearance function was assessed by the above-mentioned NR retention assay. As the results already show in Figure 2E, once again, compared to control cells, the cells likely demonstrated limited efficacy in removing the NR from the lysosomes after 48 hours of treatment with 100 µM labile iron, as evidenced by the continued existence of red puncta throughout the clearance time. Remarkably, as the clearance time passed, red puncta signals in the treated cells that were in renewed media had lessened as the clearing time passed (Figure 3G), as compared to prolonged exposure to labile iron at 48 hours (Figure 2E). From these results, we can imply that lysosome clearance function alteration by labile iron can be alleviated by renewing the fresh medium. As far as the accumulation of misfolding proteins was concerned, Th T staining was also carried out after renewing the fresh medium. Clearly, the protein aggregation signal was not observable in the control cells that were being renewed in the fresh medium; however, it was detected within the labile iron-treated cells (Figure 3H). This suggests that such a protein aggregation induced by labile iron cannot be easily cleared. Therefore, it can be said that prolonged exposure to labile iron could promote peptide or protein aggregation, which can cause cell damage and eventual death.

Next, we established a rat model with a labile iron diet to investigate whether labile iron supplementation could alter brain proteostasis and behavior. In our model, rats were given a labile iron diet (ID group) for 8 weeks (considered as the “induction phase”), and were further given a normal diet for another 8 weeks (considered as the “satellite phase”) while the other rats that were given only a normal diet (the “ND group”) were used as control (Figure 4A). This model was adapted from the established model for determining iron overload condition (43, 44), but iron source was not the same. In order to determine whether labile iron had a negative impact on behavior, we performed behavior tests (as described in Figure 4B) during the end of the induction phase and the satellite phase (weeks 13 and 14). The findings from the MWM test conducted during the induction phase indicated that the ID group had a reduced latency time (time required to locate the platform) in comparison to the ND group, particularly on day 3 of training phase (Figure 4C, top left panel). In the probe trial, there was no significant difference between ND and ID groups (Figure 4C, top left panel), suggesting that labile iron did not significantly affect spatial learning and memory in rats. In the NOR test, the results also showed that there were no significant differences between the groups in terms of the percentage of investigation time (Figure 4C, top right panel). It should be noted that both groups had a recognition index greater than 50%, indicating that the rats spent more time exploring the novel object than the familiar one during the test phase. According to the RAM test, the ID group showed an insignificant tendency to have a higher number of both working and reference memory errors. (Figure 4C, bottom lelf panel), suggesting that labile iron may have some effects on memory in rats. Whether rats given by ID could increase iron accumulation in the brain, *R*_2_* values of rat brain were determined using 1.5 tesla MRI scanner to indirectly assess iron overload condition (as illustrated in Figure 4B, right panel) (45, 46). The results showed no significant differences of *R*_2_* values between ID and ND groups (Figure 4C, bottom right panel) and the *R*_2_* values of both two groups were not as high as that observed in iron overload conditions, which usually is found in late stages of some dementia (8). Therefore, rats given the labile iron diet could not promote iron overload in rat brains. By conducting the same experiments at the end of satellite phase, similar results were observed where no significant differences in behaviors and *R*_2_* occurred between the ND and ID groups (Figure 4D). From the above results, it could be said that labile iron may not be detrimental to behavior and iron accumulation in the brain. Although the lack of significant differences may limit the interpretation of these findings, the above results suggests that labile iron can be a confounded factor on learning and memory in rats. Indeed, additional research is required to establish the underlying cause of the significantly observed changes induced by labile iron in these behavior tests.

**Figure 4.**
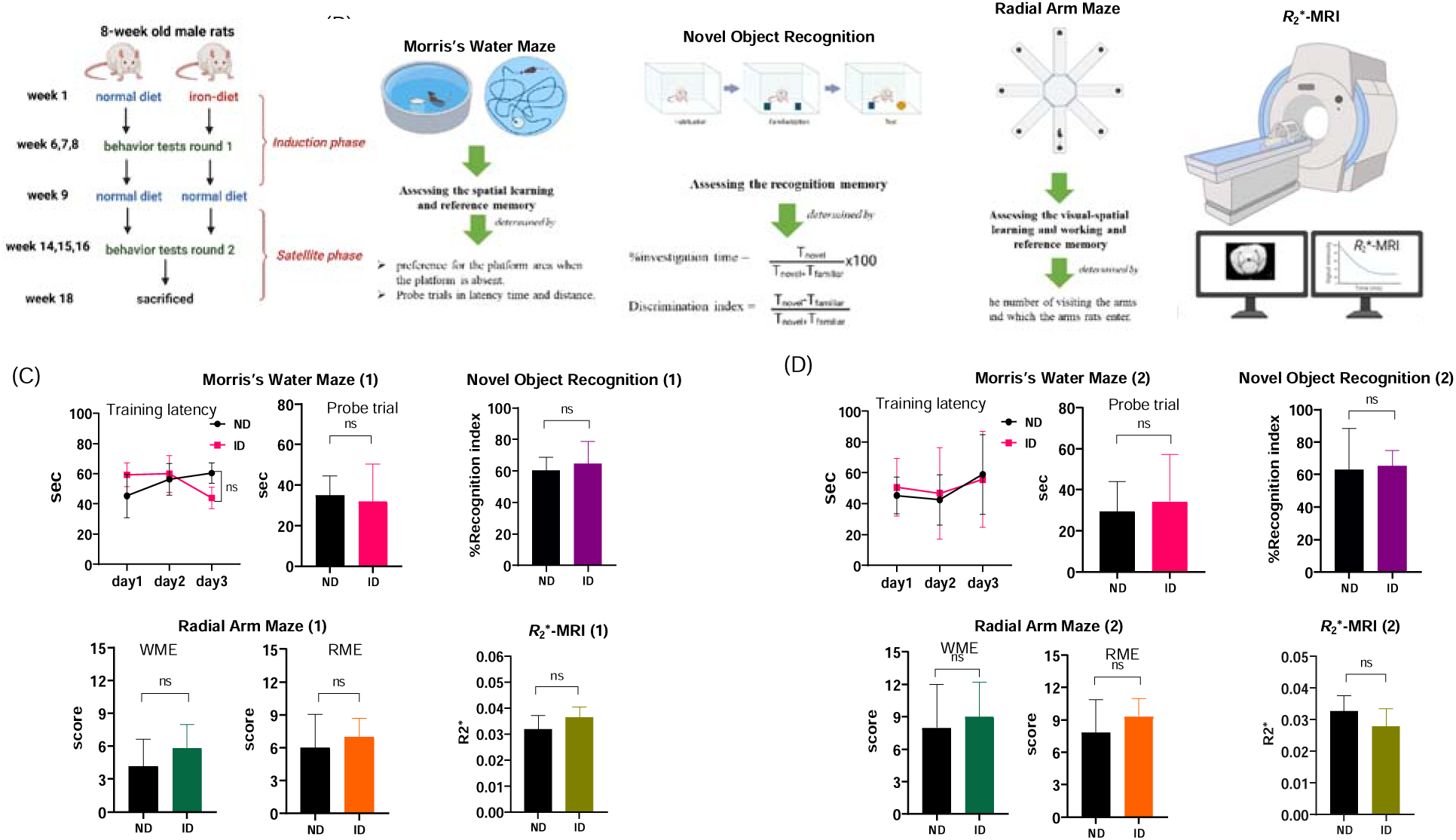
An investigation into the effect of labile iron on behavior and *R*_2_*-MRI using labile iron-diet-fed rat model. (A) Chronology of rat experiment and animal models (B) Behavioral determinations used in this study including Morris Water Maze (MWM), Radial Arm Maze (RAM), and Novel Object Recognition (NOR) examinations, as well as an *R*_2_* MRI determination, respectively. (C, D) The training latency time and probe trial in MWM test, the recognition index percentage in NOR, working memory error (WME) and reference memory error (RME) during RAM testing, and *R*_2_* value of the brain of rats in ND and ID groups during induction phase and satellite phase, respectively (Behavioral tests n=5,6, MRI n=3).

We next performed quantitative and qualitative analysis of iron accumulation in tissues. Quantification of whole brain iron levels revealed that no significant differences were observed between groups (Figure 5A). Notably, however, iron staining showed that rats in ID group could have a higher accumulation in the brain in certain regions of the brain, especially the cerebral cortex (Figure 5B). These results suggest that the labile iron supplementation could alter distribution of iron in the brain without elevating total iron level. Recently, some reports have shown that perturbed iron distribution in the cortex could lead to neurodegenerative diseases and psychiatry disorders (47). Therefore, there may be great interest to intensively study the neurobiology of labile iron in the cortex. This could pave the way to unveil a new biology of labile iron, which is implicated in novel neuronal functions.

**Figure 5.**
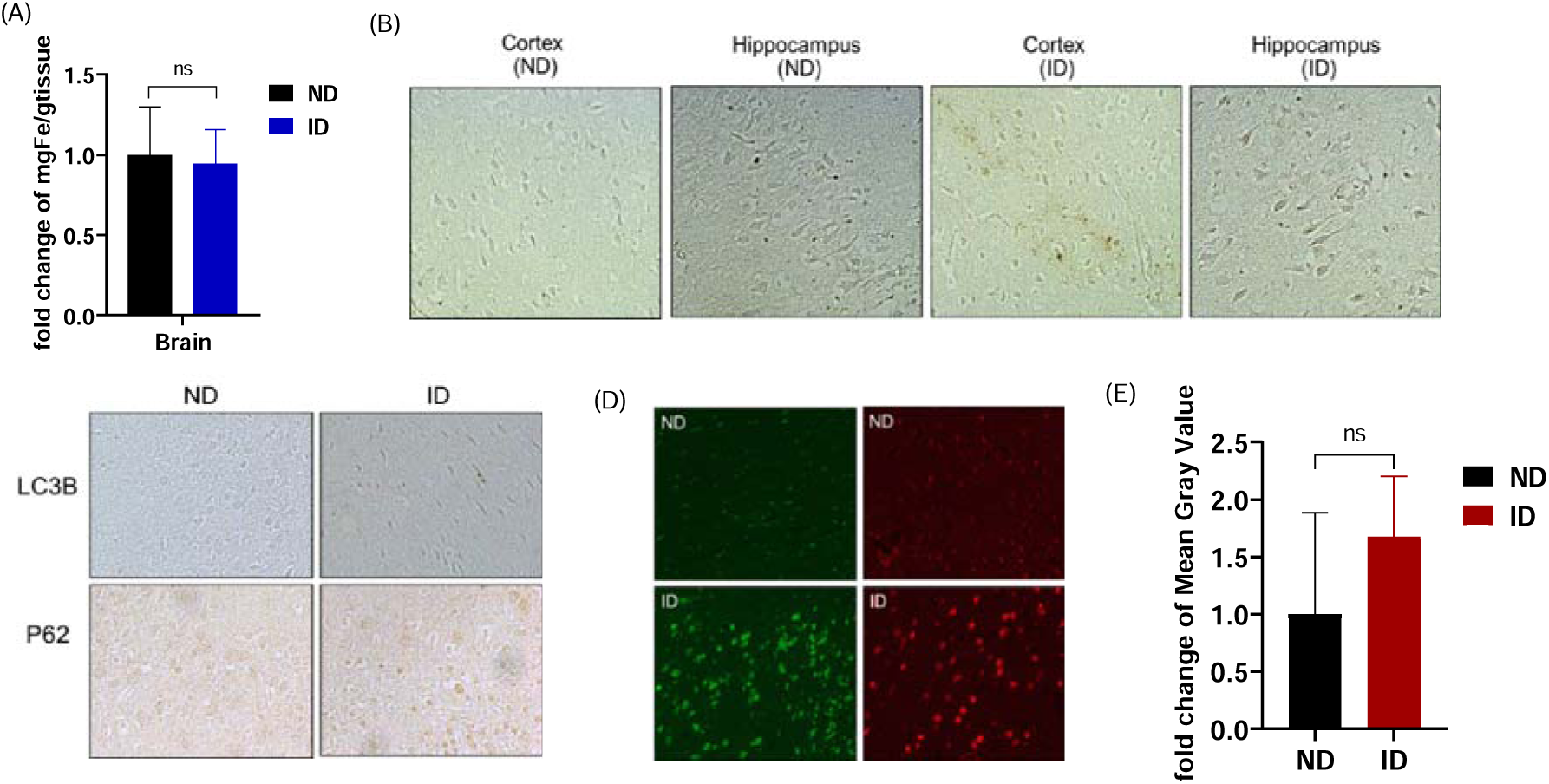
The role that labile iron plays in fostering the aggregation of proteins and its connection to autophagy in brain tissue. (A) An assessment of the iron concentration in the whole brain (n=6). (B) The determination of the iron content in the brain cortex employing Prussian’s blue staining with DAB enhancement was performed. (C) Immunohistochemistry of autophagy markers including LC3D and p62. (D) The protein aggregation in cerebral cortex were assessed by ThT staining (observed as green fluorescence), and congo red staining (observed as red fluorescence. (E) The fold change in the intensity of the mean gray value of congo red fluorescence between ND and ID (n=3).

To confirm the role of labile iron on alteration in proteostasis *in vivo*, especially in the cortex where a higher accumulation of iron was observed, an evaluation was performed. Following the sacrifice of the rats, we first determined autophagy markers in the cerebral cortex, where iron deposition was found to be observed. There was no observed difference in IHC staining of LCB expression between the ND and ID groups. However, the expression of p62 was found to be significantly higher in the cortex of rats in the ID group (Figure 5C). This finding implies that labile iron may have an impact on autophagic flux.

As far as accumulation of protein aggregation was concerned, Th T and Congo red staining were performed. As a result, the higher fluorescent intensity was observed in the cortex of rats in the ID group, compared to the ND group (Figure 5D). A semi-quantitative analysis of protein aggregate signal also suggested that the labile iron was capable of inducing accumulation of protein aggregation (Figure 5E). Overall, our findings imply that iron accumulation and its effect in the iron-diet-fed rat model were not deleterious. Nevertheless, it should be noted that even short-term exposure to labile iron may induce consequences for protein aggregation and autophagy, but this will require further study.

## Conclusions

Proteostasis and iron are not only crucial for brain functions but also are considered hallmarks of brain aging. The connection between them is still unclear. Herein, we first investigated the effect of labile iron, using a labile iron representative, on different aspects of proteostasis of neuronal cell lines. There was no doubt that much higher concentrations of labile iron resulted in increased cell death, an elevated risk of cell cycle alteration, as well as DNA damage. As far as proteostasis was concerned, a non-toxic dose of labile iron was chosen. It was found that labile iron could disturb lysosomal clearance function, autophagy, and protein translation along with a capability for protein aggregation. Studying *in vivo* using rat model with a labile iron diet, also suggests that labile iron supplementation cannot promote iron overload condition and does not have any detrimental effect on behavior tests including the MWM, RAM, and NOR, as compared to control rats. However, labile iron could alter iron transportation within the brain, resulting in higher iron deposition in certain parts of the brain such as the cerebral cortex. By staining autophagic makers (LC3B and p62) and protein aggregation in the cerebral cortex, the labile iron was found to induce p62 expression and accumulation of protein aggregation, suggesting that labile iron may induce alteration of autophagy and protein folding. Overall, our findings support the connection between proteostasis and iron, which could be a profound factor for initiation and progression of neurological disorders. Of course, much effort needs to be made to unveil the exact mechanisms for proteostasis and iron, which may serve as a valuable source of information for the treatment and prevention of neurological disorders.

## Supplementary data

Supplementary data to this article can be found online at

## Author Contributions

**Aiyarin Kittilukkana:** Conceptualization, Methodology, Investigation, Formal analysis, Writing – original draft. **Jannarong Intakhat:** Investigation. **Chalermchai Pilapong:** Conceptualization, Methodology, Supervision, Funding acquisition, Writing – review & editing

## Notes

The authors declare no competing financial interests.

## Acknowledgments

This project was funded by the National Research Council of Thailand through the RGJ-Ph.D. Program (project code: N41A650085) a Mid-Career Research Grant (project code: NRCT5-RSA63004-10). This research was partially supported by Chiang Mai University.

## Notes

### Competing Interest Statement

The authors have declared no competing interest.

